# Extracellular Hsp90α Detoxifies β-Amyloid Fibrils Through an NRF2 and Autophagy Dependent Pathway

**DOI:** 10.1101/2021.04.16.440151

**Authors:** Ayesha Murshid, Benjamin J. Lang, Thiago J. Borges, Yuka Okusha, Sachin P. Doshi, Suraya Yasmine, Joanne Clark-Matott, Reeham Choudhury, Lay-Hong Ang, Maya Woodbury, Tsuneya Ikezu, Stuart K. Calderwood

## Abstract

We have investigated the role of <u>e</u>xtracellular <u>H</u>eat <u>s</u>hock <u>p</u>rotein <u>90</u> <u>alpha</u> (eHsp90α) in conferring protection of neuronal cells against fibrillary amyloid beta (f-Aβ_1-42_) toxicity mediated by microglial cells. Formation of f-Aβ_1-42_ plaques leads to neurotoxic inflammation, a critical pathological feature of Alzheimer’s Disease. We observed increased uptake and clearance of internalized f-Aβ_1-42_ by microglial cells treated with eHsp90α, an effect associated with activation of NRF2 (NF-E2-related factor 2) - mediated autophagy. eHsp90α thus mitigated the neuronal toxicity of f-Aβ_1-42_-activated microglia. In addition, eHsp90α facilitated f-Aβ_1-42_ engulfment by microglial cells *in vitro*. In summary, eHsp90α triggers NRF2-mediated autophagy in microglia and thus protects against the neurotoxic effects of f-Aβ_1-42_.

## Introduction

Intracellular HSPs are stress proteins that mediate cell survival during the heat shock response (HSR) through maintenance of protein homeostasis (proteostasis) [1-3]. Such HSPs support survival by guiding protein folding and modulating protein degradation, thus protecting against proteotoxic stresses such as heat shock. However, in addition to the intrinsic HSR, the response to proteotoxicity has been shown to be transcellular in nature and protein stress at one site can modulate the response in cells at distant sites. It has become clear now that HSPs such as Hsp90 are abundantly secreted into the extracellular microenvironment and such extracellular HSPs (HSPe) may be important components of the transcellular HSR. HSPs released in free form or in extracellular vesicles may be able to interact with distant cells to increase their capacity for proteostasis. In addition to boosting cellular chaperone levels, HSPs may trigger signaling pathways that influence cell phenotype on encountering receptors in target cells. In mononuclear phagocytes, the cells under study here, HSPs have been shown to bind to the scavenger receptors LOX-1 and SREC-1. Ligand-associated LOX-1 is known to activate the factor NFκB through signaling pathways involving generation of reactive oxygen species (ROS).

In the current study, we have examined the role of extracellular Hsp90 (Hsp90e) in the responses of microglia, brain resident mononuclear phagocytes that are the principal immune cells in the central nervous system (CNS). We have examined the role of Hsp90e in survival respoonses of micrglia as well as in their potential role in neuroinflammation during beta amyloid disorder.

These activities may have therapeutic relevance for neurodegenerative diseases such as Alzheimer’s Disease (AD) and increasing the expression of HSPs, particularly Hsp90, has been suggested as an approach to manage the morbidity of AD [8-10]. AD is a progressive neurodegenerative disease characterized by loss of neuronal cells, accumulation of f-Ab aggregates and intracellular hyperphosphorylated tau [11]. Ab peptides are generated in cells by digestion of the amyloid precursor protein (APP) protein in membranes by secretases. The peptides vary in their abilities to form amyloid fibrils with Ab_1-42_ highly amyloidogenic compared to other products such as Ab_1-40_ which has little known toxicity [12]. Microglial cells, the resident mononuclear phagocytes of the brain, are considered vital in removing the toxic f-Ab_1-42_ aggregates from the extracellular milieu of neuronal tissues [13]. Following uptake by microglia, f-Ab_1-42_ aggregates are typically transported to the endolysosomal pathway for degradation. However, this process may initiate microglia to adopt an inflammatory phenotype that is toxic to surrounding neuronal cells and is ultimately a key component of AD pathogenesis [14, 15]. These activities of microglia upon f-Ab_1-42_ metabolism therefore constitute a “double edged sword”; on one hand reducing levels of extracellular f-Ab_1-42_ and on the other initiating a neurotoxic inflammatory environment.

We have investigated whether exogenous delivery of Hsp90α could ameliorate the potentially malign influence of exogenous f-Ab fibrils (f-Aβ_1-42_) upon microglial cells and subsequent neuronal toxicity. Indeed, eHsp90α played a significant role in protecting neuronal cells from f-Aβ_1-42_ aggregates in the presence of microglia. The protective activities of eHsp90α appeared to be multifaceted. Of particular interest, addition of Hsp90α re-directed internalized f-Aβ_1-42_ to autophagosomes, an effect likely facilitated by activation of the NRF2 detoxification pathway and was associated with reduced production of the nitric oxide (NO) by-product nitrite. In addition, eHsp90α facilitated microglial f-Aβ_1-42_ uptake. We have therefore demonstrated a cytoprotective signaling pathway activated by eHsp90α that is effective in ameliorating the toxic effects of f-Aβ_1-42_.

## Materials and Methods

### Cell Culture and siRNA construct transfection

Most studies were carried out in BV2 cells, a line of immortalized mouse microglia [16]. Some confirmatory studies used EOC2, an immortalized microglial cell line derived from the brain of an apparently normal 10-day-old mouse [17]. Cells were cultured in Dulbecco’s Modified Eagles Medium (DMEM) F12 containing 10% FBS and 1% L-glutamine. BV-2 cells were maintained in DMEM supplemented by 10% HI FBS and Penicillin-Streptomycin (1000 units/ml), non-essential amino acids, HEPES, monocyte colony stimulating factor (M-CSF, 20ng/mL, R&D Systems). EOC2 cultures were maintained in DMEM media supplemented with 10% HI FBS, LADMAC media (20%) and 2ml L-glutamine. HT22 cells were sourced from INSERT and maintained in MEDIA-X. Primary murine microglia were cultured and maintained according to [18]. All cell cultures were maintained in a 5% CO_2_ humidified incubator at 37°C.

Microglial cells are plated into pre-coated inserts, which fit into wells of 24-well plates. The plating surface of the insert consists of a porous nylon mesh (3.0 uM), which allows soluble factors secreted by microglia to become a part of the shared neuron (HT22) -microglia environment.

### Animals, Chemicals and antibodies

Pregnant CD-1 mice used to prepare primary microglia were purchased from The Jackson Laboratory. Mice were housed under standard Laboratory conditions (23±1°C, 55±5% humidity) and had continuous access to drinking water and food. Neonatal murine microglia were isolated from P0 CD-1 pups using CD11b microbeads (Ca #130-093-634, Miltenyi Biotec) and their purity was assessed by immunocytochemistry of myeloid cell markers (Iba-1 and CD11b) according to the published method [19]. The experiments were performed in accordance with *the Guidelines for the Institutional Animal Care and Use of laboratory animals of the Boston University School of Medicine (IACUC #15178)*.

Mouse Aβ_1-42_ and Aβ_1-40_, FITC tagged mouse Aβ_1-42_ and control peptides were purchased from American Peptides and AnaSpec. Recombinant full-length human Hsp90α was expressed by baculovirus in Sf9 insect cells using a C-terminal His tag vector, purified by metal affinity chromatography and thus prepared free of endotoxin contamination [7]. Anti-rabbit Hmox-1 antibodies and anti-β-actin mouse monoclonal antibodies were from Sigma-Aldrich, Anti rabbit NRF2 and Anti-rabbit phospho-NRF2 were from Abcam. β-tubulin was from Abcam. MAP1-LC3B polyclonal antibodies were from Sigma-Aldrich and Cell signaling Technology Inc. Sqstm1 (p62) antibodies were from Cell Signaling Technology Inc. Murine Macrophage colony stimulating factor (M-CSF) was sourced from R&D Systems.

### Preparation of Aβ fibrils

Aβ_1-42_ was dissolved in DMSO (stock 500 µM) at room temperature and stored at -20°C. To this Aβ aliquot, we added 10 mM HCl at RT, diluting to a final concentration of 100 μM of fAβ_1-42_. We mixed by vortex for 15 s, transferred the solution to 37°C and incubated for 24 h. The fAβ_1-42_ solution was then incubated for 24 h at 37°C.

### Western Analysis

Cells were washed extensively in ice-cold phosphate-buffered saline, pH 7.4 (PBS) and protein lysates prepared in RIPA lysis buffer containing 1% NP-40, 0.5% sodium deoxycholate and 0.1% SDS in the presence of a protease and phosphatase inhibitor cocktails. Protein samples (20μg) were then subjected to 4–15% gradient gel SDS-PAGE using the standard *Cold Spring Harbor Laboratory* protocol and transferred electrophoretically to PVDF membranes. Filters were then blocked in 5% bovine serum albumin (BSA) and probed for 2h with either anti-MAPI-LC3B (used at 1:500 dilution), anti-β-actin (1:5000), anti-pNRF2 (S40, 1:4000), anti-NRF2 (1:200), anti HmoxI (1:500), anti p62 (1:1000), or anti GAPDH (1:300) and antibody-antigen complexes visualized as described [20] using chemiluminescent ECL reagents. The LC3I and LC3II isoforms were distinguished by differential electrophoretic mobility. Although higher in molecular weight, LC3-II migrates more rapidly in the electrophoretic field due to its modification with phosphatidylethanolamine.

For re-probing blots with multiple antibodies, they were stripped overnight in buffer containing 1.5% glycine, 0.1% SDS, and 1% Tween 20 at pH 2.2, re-blocked with 5% BSA and reprobed with the next antibody.

### Immunofluorescence and confocal microscopy

At the indicated time points, BV-2 and EOC2 cells were washed in ice-cold PBS, pH 7.4, fixed with 4% para formaldehyde at room temperature, and then permeabilized with 0.1% Triton X-100. Cells were then blocked with 3% normal goat serum for 1 h at room temperature. Iba1 (Abcam,cat. no. ab120481) and fluorophore-tagged secondary antibodies were used to fluorescently stain the fixed cells and nuclei were stained with DAPI. Coverslips were mounted with *Prolong Gold* medium. Slides were scanned using a Zeiss LSM 810 confocal microscope with Zen software with the respective, appropriate filter sets as previously described [7]. Neurite growth in Fig 1E was measured using Imaje J software. Colocalization was quantified in Zen Software and with Adobe Photoshop software with the help of Channel on/off.

**Figure 1.**
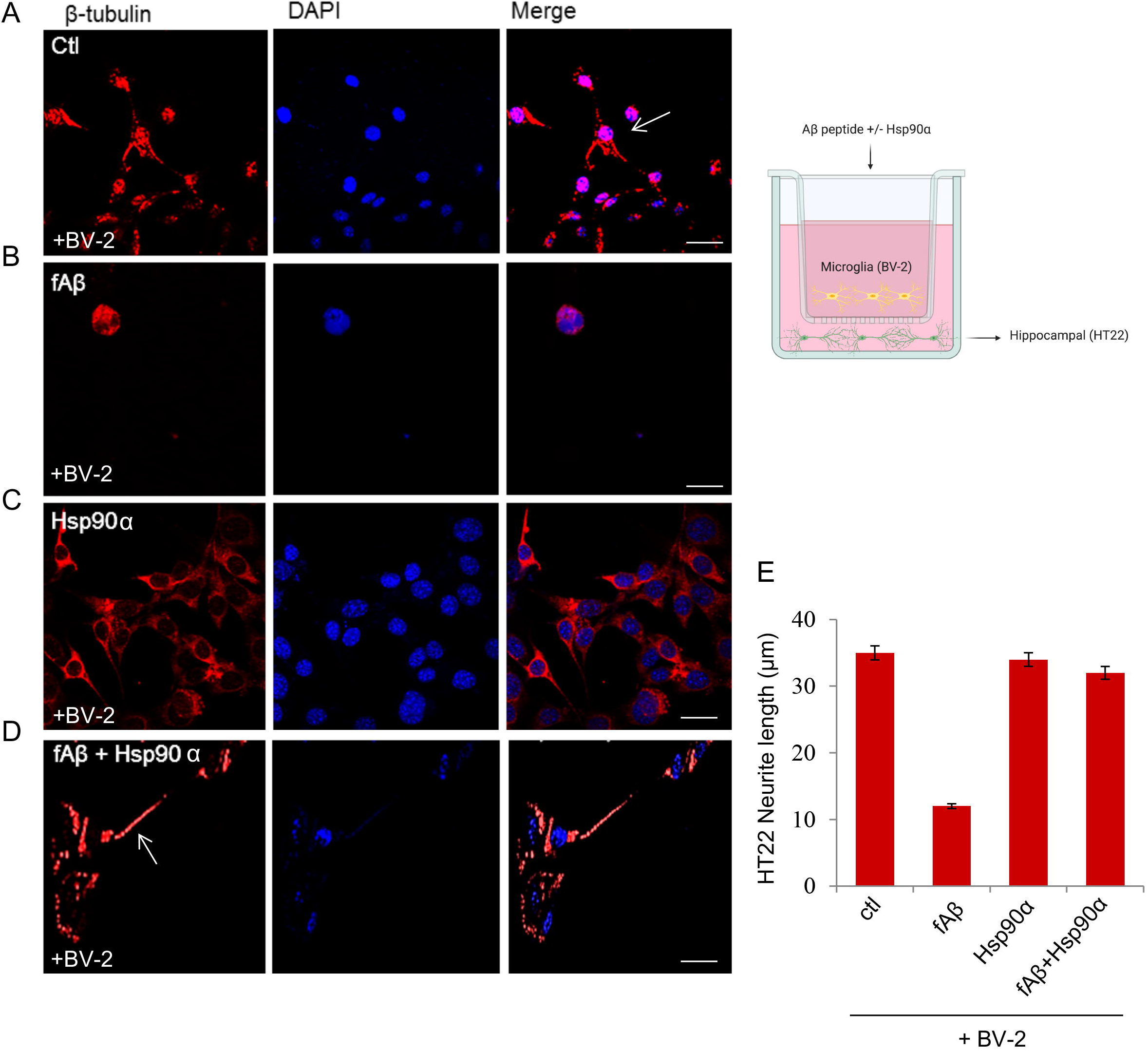
eHsp90α mitigates Fibrillar Amyloid β induced Neurotoxicity. (A-D). HT22 hippocampal neuronal cells were grown on coverslips in the bottom layer of a transwell culture dish. BV-2 cells were then added to the top layer of the transwell, and incubated with: (A) no ligand, (B) fibrillar f-Aβ_1-42_ (2µM), (C) Hsp90α (10µg/ml) and (D) f-Aβ_1-42_ + Hsp90α (D) for 72 h. After the 72 h incubation, HT22 cells from the bottom wells were then fixed with 4% para formaldehyde and then permeabilized with 0.1% Triton X-100 before staining with anti β-tubulin antibodies. Stained cells on coverslips were then examined by confocal microscopy. (E) β-tubulin stained neurite outgrowth was measured using image J. scale bar = 5µm A total of 100 cells were counted in each sample. Cartoon created with *BioRender.com*. Experiments were repeated three times with similar results.

### Quantification of Nitric oxide production

Nitric Oxide (NO) release from cells was measured (using the manufacturer’s protocol) in cells incubated with or without f-AB_1-42_ and in other control samples using a Nitric Oxide (total) detection kit, Cat# ADI-917-020.

### RNA isolation and RT-qPCR

Total RNA was isolated using the RNeasy Mini kit (*Qiagen*), including on-column DNase digestion to eliminate DNA (Rnase-Free DNase Set, Qiagen). RNA quantification was then performed using the Spectrophotometer ND-1000 (NanoDrop). RNA was reverse-transcribed using the iScript cDNA Synthesis Kit (Bio-Rad) or an Applied Biosystems kit. cDNA (20 ng) was amplified using the following Taqman Gene Expression Assays (ThermoFisher Scientific): cDNA (20 ng) was amplified using the following Taqman Gene Expression Assays (ThermoFisher Scientific): *Nfe2l2* (Mm00477784_m1), *Hmox1* (Mm00516005_m1), *Nqo1* (Mm01253561_m1), *Sqstm1* (Mm01070495_m1) and *18s* (Mm03928990_g1). All qPCR reactions were performed in a *StepOne Plus Real-Time PCR System* (Applied Biosystems). The relative mRNA levels were calculated using the comparative Ct method, with 18S as the internal control.

### Statistical analysis

We determined differences between two specific points using the Student’s t-test. To determine differences between three or more groups, we used the one-way analysis of variance (ANOVA) test, with Tukey post hoc tests. The level of significance was set at p<0.05. All analyses were performed using the software Prism 6 (GraphPad Software Inc.).

## Results

### Extracellular Hsp90α mitigates Fibrillar Amyloid beta-induced Neurotoxicity *in vitro*

To test whether eHsp90α could protect neurons from the inflammatory toxicity of f-Aβ_1-42_ -activated microglia we co-treated murine microglial BV-2 and EOC2 cells with purified Hsp90α and/or freshly prepared f-Aβ_1-42_ and assessed the viability of adjacent neuronal HT22 cells potentially exposed to secreted microglial products via transwell culture preparations. BV-2 cells pre-incubated with FITC-fAβ_1-42_ mediated toxicity towards the distant neuronal HT22 cells as indicated by extensive loss of microtubule-containing processes, a morphological measure of neuron cell viability [21] (Fig. 1A, B). Within 72 h of f-Aβ_1-42_ incubation, many of the neuronal cells (HT22) had lost their elongated processes (Fig. 1B). In contrast, when HT22 cells were co-cultured with BV-2 cells that had been treated with both f-Aβ_1-42_ and eHsp90α in the top well of the transwell culture dish, the majority of the HT22 survived with neurite lengths comparable to those in the non-treated control, suggesting protection by the chaperone (Fig. 1D). eHsp90α treatment alone did not significantly impact neurite outgrowth (Fig. 1C). The extent of neurite outgrowth was quantified and is shown in Fig. 1E. Similar effects of exposure to f-Aβ_1-42_ without or with eHsp90α were also found upon co-culture of HT22 neuronal cells with EOC2 microglial cells (Suppl. Fig. 1).

### Nitric Oxide secretion by microglia is reduced by eHsp90α

It was recently reported that microglia can express high levels of inducible nitric oxide synthase (iNOS) upon internalization of f-Aβ_1-42_ and such elevated iNOS leads to an increase in NO secretion [22]. Such high levels of NO become highly toxic after reaction with oxygen to form peroxynitrate (ONOO^—^), conferring lethal DNA damage to adjacent cells. Both increased iNOS expression and increased NO production have been shown to be contributing factors in fAβ-induced neurotoxicity [22, 23]. We therefore measured NO secretion by BV-2 cells treated with f-Aβ_1-42_, eHsp90α or co-treated with eHsp90α and f-Aβ_1-42_ and observed that while f-Aβ_1-42_ increased levels of NO secretion by approximately 4-fold, samples co-treated with eHsp90α and f-Aβ_1-42_ produced comparatively lower NO levels (Fig. 2A). These data suggested that one potential mechanism by which exposure to eHsp90α might reduce f-Aβ_1-42_-associated neurotoxicity could be induction of the anti-oxidant response pathway through NRF2 activation, with increased induction of its anti-oxidative gene targets and subsequent reduction in NO levels [24]. To further test this hypothesis, *Nfe2l2* mRNA levels encoding the NRF2 protein were knocked down in BV-2 cells using siRNA and cells were then incubated with f-Aβ_1-42_, eHsp90α or co-treated with eHsp90α and f-Aβ_1-42_. While exposure to eHsp90α reduced NO accumulation by f-Aβ_1-42_ in the scrambled control cells, this sparing effect was attenuated in the *Nfe2l2* siRNA sample, supporting a role for NRF2 in the protective properties of eHsp90α against f-Aβ_1-42_-associated NO production (Fig. 2B).

**Figure 2.**
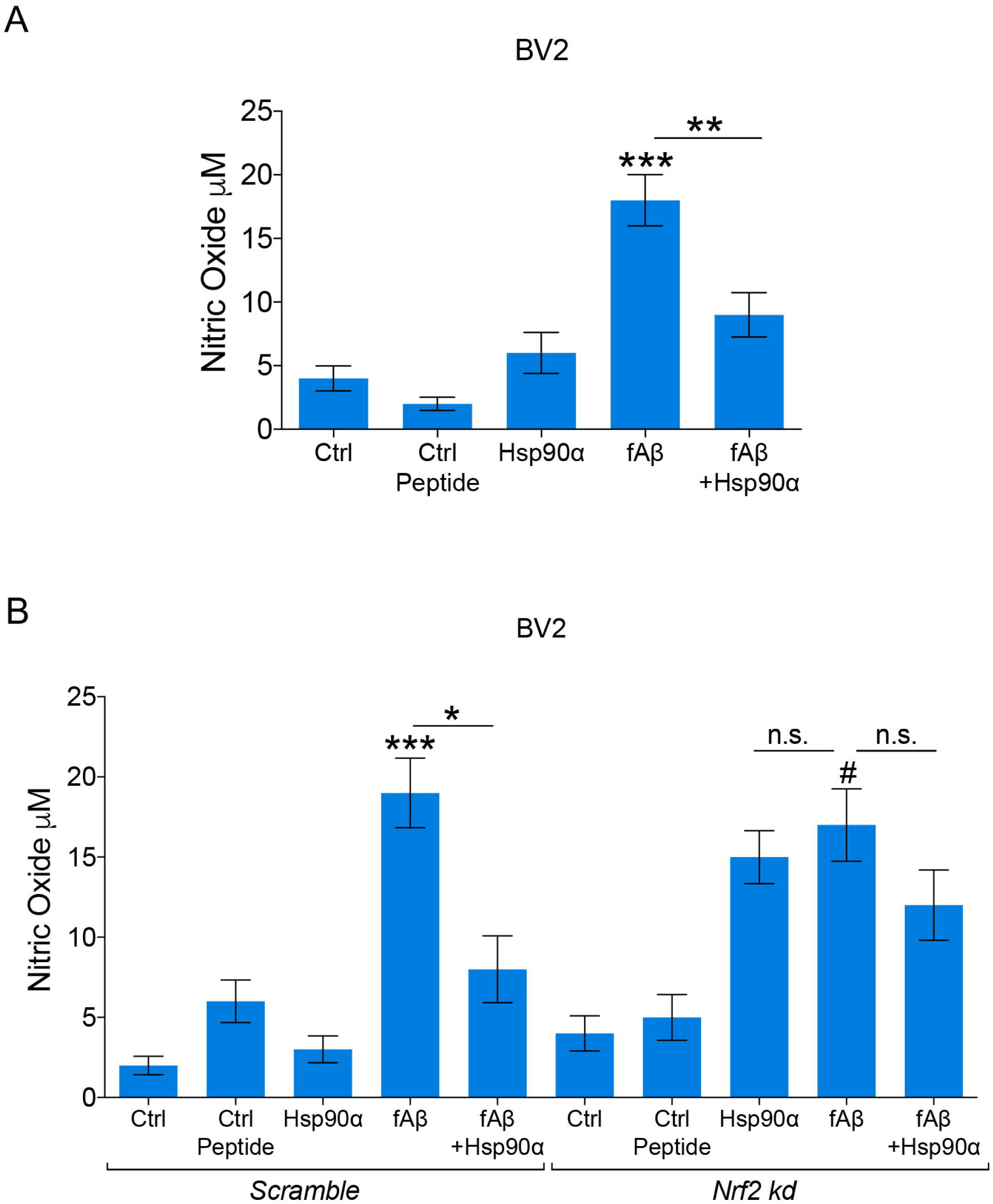
Nitric Oxide secretion induced by microglial exposure to fibrillar Aβ is reduced by eHsp90α. (A) BV-2 cells were incubated with the indicated ligands or with vehicle (ctl) for 4-6 h. Control peptide was Aβ_1-40_. NO secretion to the medium was then quantitated using a Pierce assay kit, according to manufacturer’s protocol. (B) BV-2 cells were transfected with scrambled control (*scr)* RNA or *Nfe2l2* siRNA for 72 h. Cells were then incubated with indicated ligands or not (ctl) for 6 h and NO secretion was measured as above. Experiments were repeated 3 times with similar results.

### eHsp90α activates the NRF2-antioxidant response element signaling pathway in BV-2 cells

Activation of NRF2 and its antioxidant gene products such as Hmox1 (Heme Oxygenase 1), NQO1 (NAD [P] H:quinone oxidoreductase, Prdx1 (peroxiredoxin), leads to protection of cells from inflammatory damage through ROS (reactive oxygen species) [25].We therefore asked whether Hsp90α could induce NRF2 and its antioxidant gene products and thus deter microglia from f-Aβ_1-42_-mediated toxicity. To test this possibility, we incubated BV-2 cells with f-Aβ_1-42_, eHsp90α or eHsp90α + f-Aβ_1-42_ and assayed for indicators of altered NRF2 activity. We first observed increases in mRNA levels of *Nfe2l2* in cells treated with eHsp90α + f-Aβ_1-42_ or eHsp90α alone, but only a mild effect of f-Aβ_1-42_ alone (Fig. 3A). We also observed upregulation of known NRF2 regulated genes, *Hmox1* and *Nqo1* mRNA in microglia incubated with eHsp90α (Fig. 3B, C).

**Figure 3.**
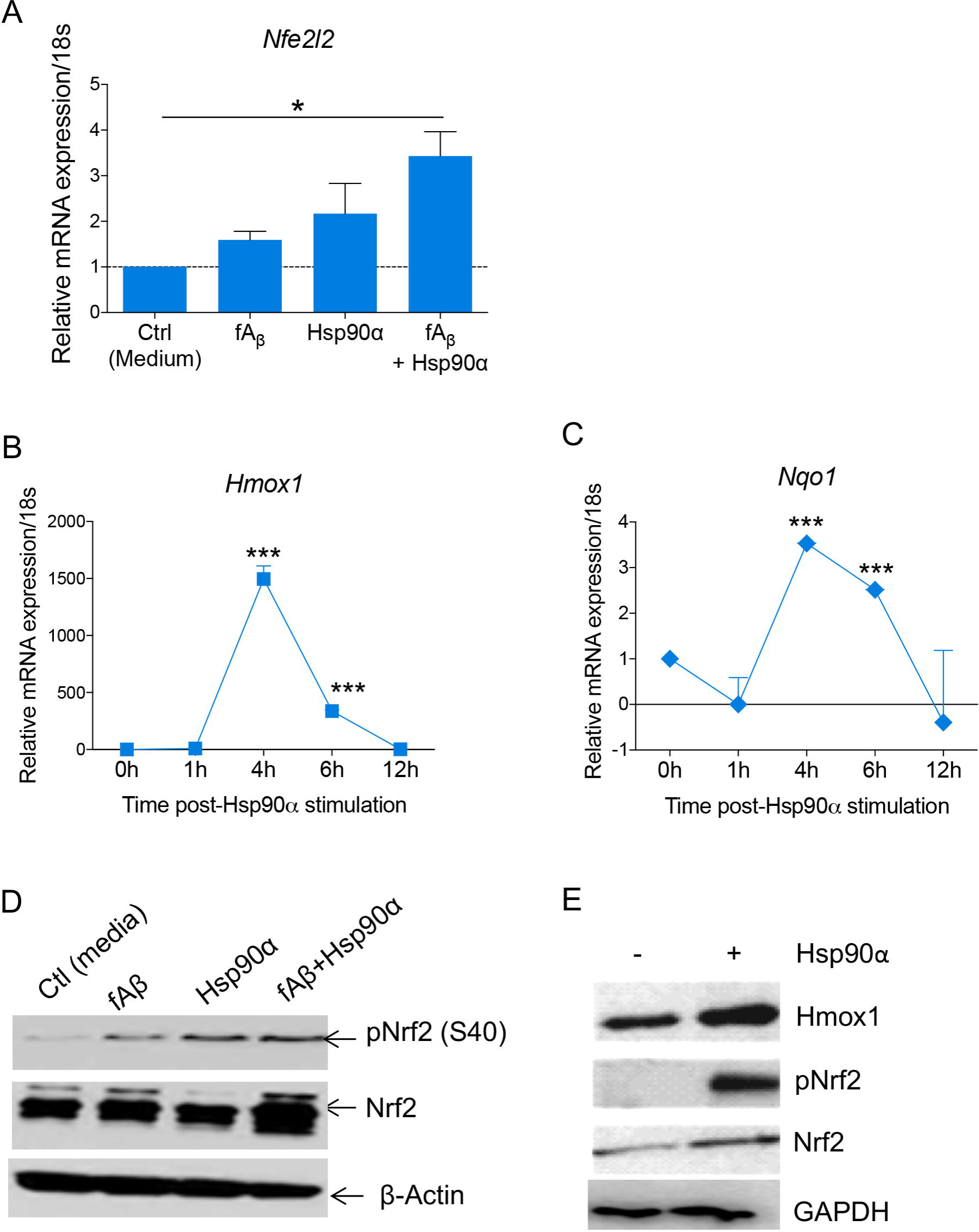
eHsp90α treatment activates the NRF2-antioxidant response element signaling pathway in BV-2 cells. (A-C) BV-2 cells were incubated with eHsp90α for the indicated times and then total RNA was extracted and assayed by RT-qPCR. Relative *Nfe2l2* (A), *Hmox1* (B) and *Nqo1* (C) mRNA expression was quantified and normalized to 18S and represented as fold-change to control or 0 h timepoint. (D, E) BV-2 cells were incubated with Aβ, eHsp90α or eHsp90α plus Aβ for 4 h. Total cell lysates were collected, separated using SDS-PAGE and analyzed by immunoblot. In D, the filters were probed with anti-pNRF2 antibodies then stripped and probed sequentially for NRF2, and β-actin (loading control). In E, filters were probed after immunoblot with anti-Hmox1 antibodies then stripped and probed sequentially for pNRF2, NRF2, and GAPDH (loading control). Experiments were repeated 3 times with similar results.

We next demonstrated an increase in the active, modified form of NRF2, phospho-NRF2 (S40) and total levels of NRF2 protein in BV-2 cells incubated with both f-Aβ_1-42_ and eHsp90α (Fig. 3D). Hmox1 protein expression was also increased in cells incubated with eHsp90α (Fig. 3E).

### eHsp90α increases the uptake and clearance of f-Aβ_1-42_ by microglial cells

Next, we tested the hypothesis that addition of Hsp90α might produce additional beneficial effects by increasing phagocytosis of f-Aβ_1-42_, thus removing some of the toxic aggregates from the medium. We therefore examined the effect of eHsp90α on uptake and accumulation of fluorescence-labeled f-Aβ_1-42_ in BV-2 and primary cultured murine microglia. Addition of eHsp90α at 10µg/ml markedly increased uptake of 2.5µM of f-Aβ_1-42_ in BV-2 (Fig. 4A-B), primary microglia (Fig. 4C-D) and EOC2 cells (Suppl. Fig. 2). Primary microglia were treated with Alexa-555-labelled eHsp90α (Fig. 4C-D). Co-localization of Alexa-555-Hsp90α and FITC-f-Aβ_1-42_ after 2 h incubation with primary microglia was observed suggesting that eHsp90α may facilitate f-Aβ_1-42_ uptake (Fig. 4D). We used anti-Iba1 antibody to mark the microglial population in primary culture. Iba1 is a microglia-macrophage specific calcium-binding protein.

**Figure 4.**
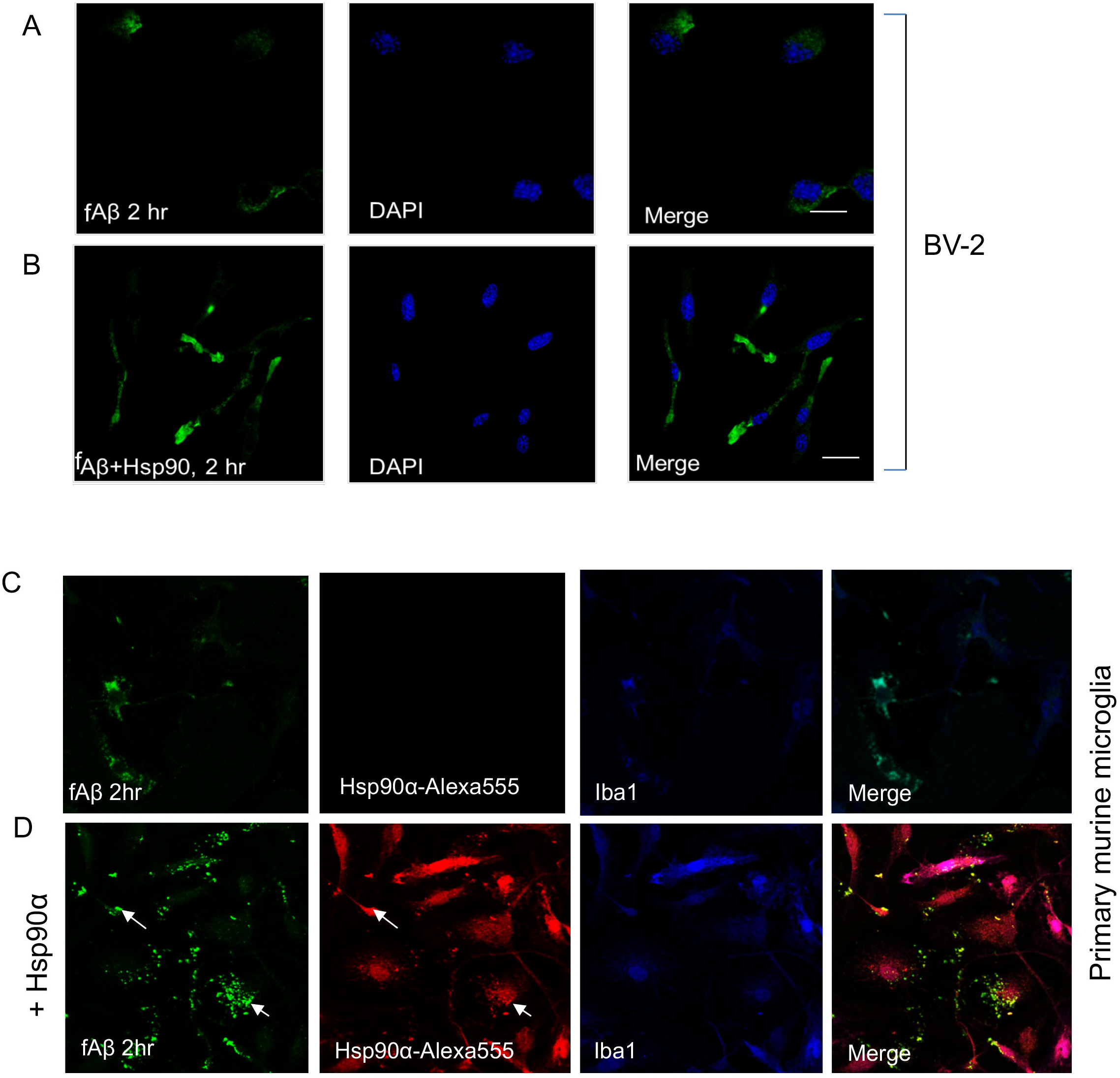
eHsp90α increases uptake of FITC-f-Aβ_1-42_ by microglia. (A, B) BV-2 cells were incubated with 2µM FITC-fAβ (green) for 2 h. Cells were then fixed with 4% paraformaldehyde, permeabilized (0.1% Triton X100) and stained with DAPI (blue). FITC-f-Aβ_1-42_ was detected by its intrinsic fluorescence. scale bar = (5 µm). (C, D) Primary microglia were incubated with 2µM FITC-fAβ (green) and ± Alexa555-Hsp90α (red) for 2 h. Cells were then fixed, permeabilized, stained with DAPI (blue). FITC-f-Aβ_1-42_ and Alexa555-Hsp90α were detected by their intrinsic fluorescence. Experiments were repeated twice, and 100 cells were counted from each sample for each experiment, scale bar = (X).

To determine the influence of eHsp90α on the fate of internalized FITC labeled f-Aβ_1-42_ after internalization in microglia, we incubated BV-2 cells (Fig. 5A-B) and EOC2 cells (Suppl. Fig. 3) with FITC-f-Aβ_1-42_ for 19 and24 h at 37°C, respectively. Samples co-treated with eHsp90α had a faster rate of intracellular FITC depletion potentially suggesting higher levels of f-Aβ_1-42_ degradation. An increased rate of cytosolic f-Aβ_1-42_ loss was also observed in the early periods upon addition of eHsp90α within 19 h of treatment (Fig. 5. Increased f-Aβ_1-42_ uptake was observed within 9.5 h in the eHsp90α co-treated sample compared to f-Aβ_1-42_ alone. This pattern was consistent with the effects observed in EOC2 cells, where higher levels of f-Aβ_1-42_ were observed within 2 h of treatment (Suppl. Fig. 2), yet at 24 h the f-Aβ_1-42_ FITC signal was observed to be reduced in the experimental group treated with both f-Aβ_1-42_ and eHsp90α compared to f-Aβ_1-42_ alone (Suppl. Fig. 3). When considered together the data suggested that, similar to the BV-2 cells, the EOC2 cells also appeared to exhibit faster rates of f-Aβ_1-42_ uptake and metabolism when co-treated with eHsp90α.

**Figure 5.**
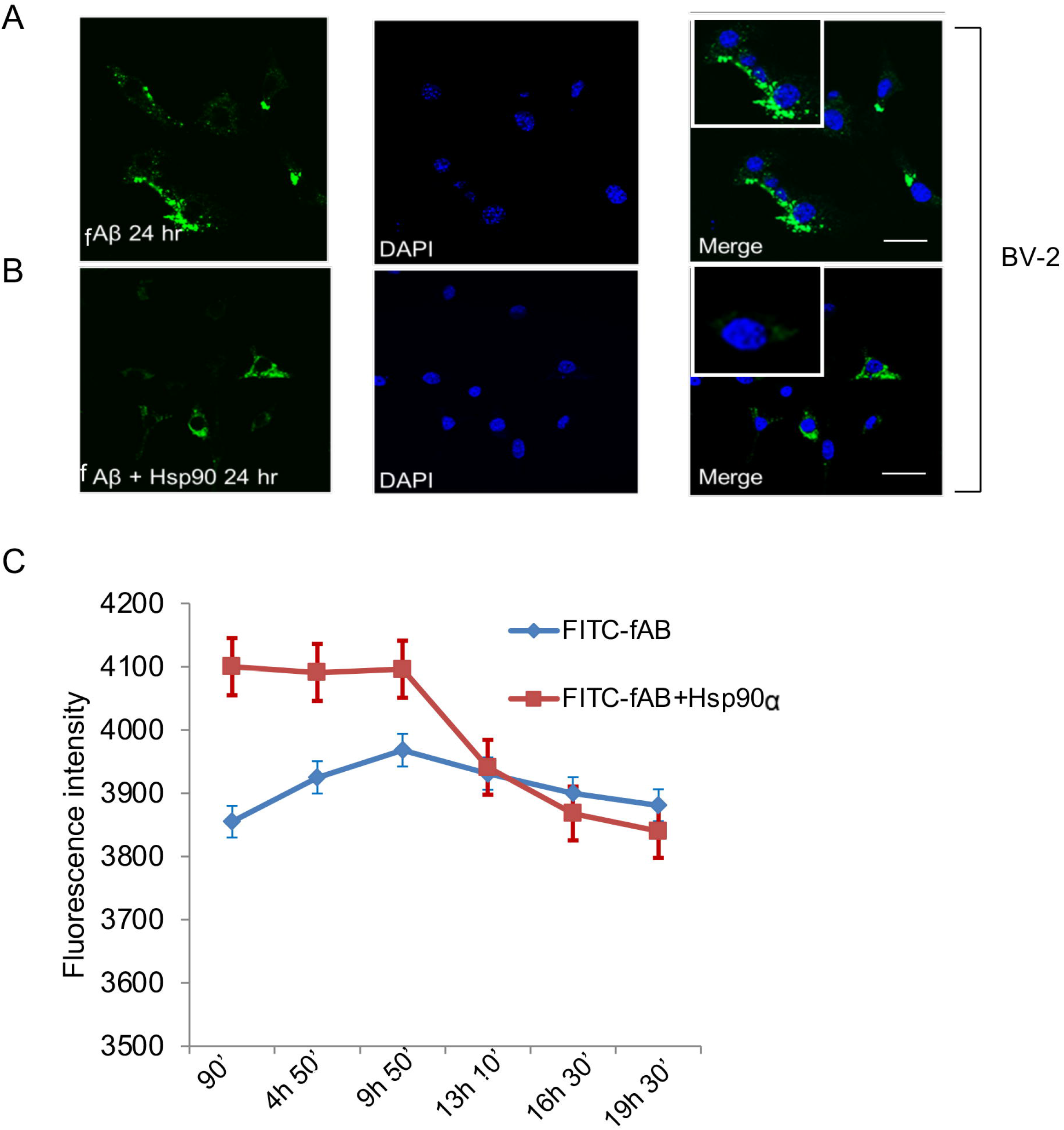
Altered clearance of FITC-fAβ_1-42_ fibrils in microglia by eHsp90α. (A, B). BV-2 cells were incubated with FITC-f-Aβ_1-42_ (green, 2µM) ± Hsp90α for 24 h. Cells were then fixed (4% para formaldehyde), permeabilized (0.1% Triton X-100) and stained with DAPI (blue) prior to confocal microscopy. (C) BV-2 cells were incubated with 2µM FITC-f-Aβ_1-42_ and/or 10µg/ml of eHsp90 for up to 19.5 h. Cells were then fixed, permeabilized and stained with DAPI (blue) as in A and B prior to determination of fluorescence intensity. Scale bar = X. Experiments were repeated 3 times, reproducibly.

### eHsp90α promotes autophagosome-mediated clearance of f-Aβ_1-42_ by microglial cells

To further investigate mechanisms through which eHsp90α might modulate the intracellular fate of f-Aβ_1-42_, we examined the intracellular localization of the fibrils after uptake and asked how this was impacted by co-treatment with eHsp90α. It has been previously shown that f-Aβ_1-42_ can be degraded by the autophagy pathway in some circumstances [26]. We found that after uptake, f-Aβ_1-42_ was localized to regions containing both the lysosomal marker LAMP1 (Fig. 6A and E) and the autophagosome marker MAP1-LC3 (Fig. 6B and E) in BV-2 cells after 2 h of treatment with FITC-f-Aβ_1-42_. However, upon co-treatment with eHsp90α the proportion of cells that exhibited co-localization of FITC-f-Aβ_1-42_ with LAMP1 was decreased (Fig. 6C and E), while cells with colocalization of f-Aβ_1-42_ and autophagosomal MAP1-LC3 marker was increased, by at least 3-fold (Fig. 6D and E). These experiments therefore suggested that eHsp90α redirects the fibrils towards the autophagosomal compartment.

**Figure 6.**
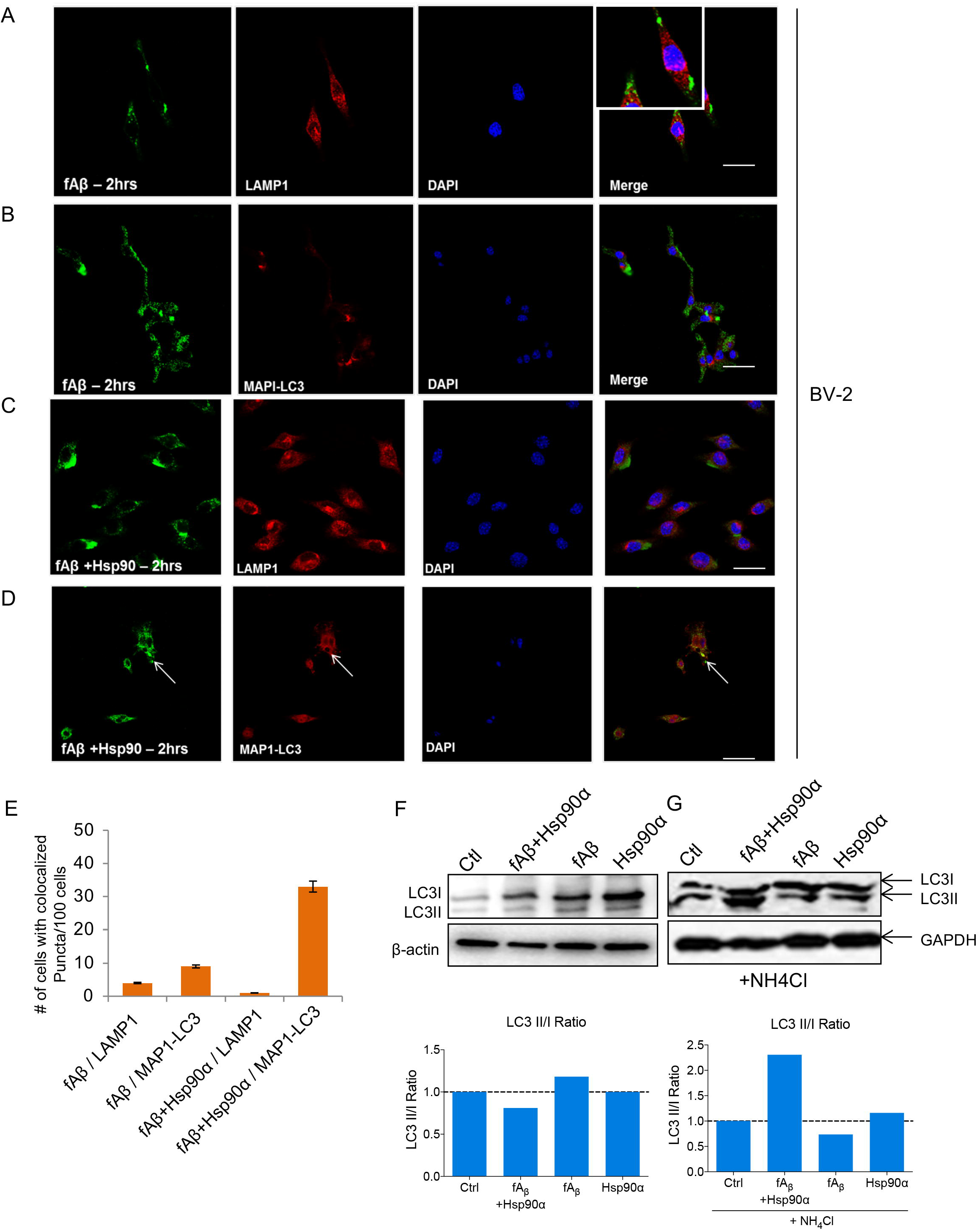
FITC-fAβ_1-42_ fibrils are localized in MAP1-LC3B marked autophagosomes in the presence of eHsp90α. (A) BV-2 cells were incubated with FITC-fAβ (2µM, green) for 2 h. Cells were then fixed, permeabilized and stained with anti-LAMP1 antibodies (red) and DAPI (blue) prior to confocal microscopy. (B) BV-2 cells were incubated with FITC-fAβ (2µM, green) as in A. Cells were then fixed, permeabilized and probed by immunofluorescence with MAPI-LC3B antibodies (red) and DAPI (blue). (C) BV-2 cells were incubated with FITC-fAβ (2µM, green) and eHsp90α (10µg/ml) for 2 h. Cells were then processed as in A, B and then stained with anti LAMP1 antibodies (red) and DAPI (blue). (D) BV-2 cells were incubated with FITC-fAβ and eHsp90α as in A and then processed as in A. Cells were probed by immunofluorescence with anti-MAP1-LC3B ab (red) and DAPI (blue). MAP1-LC3B was thus detected by secondary fluorescent antibodies and FITC-f-Aβ_1-42_ by its intrinsic fluorescence. (E) Quantitation of the relative levels of colocalized FITC-f-Aβ_1-42_ and either LAMP1 or LC3 by fluorescence intensity. (F) BV-2 cells were treated with f-Aβ_1-42_ (2µM) ± eHsp90α (10µg/ml) or vehicle for 2 h. Cells were then lysed and lysates containing equal amounts of protein were analyzed by SDS-PAGE priors to electro-transfer and immunoblot with MAPI-LC3BI antibodies. The PVDF membrane was stripped and blotted with anti-GAPDH antibodies as loading control. (G) BV-2 cells were pretreated with NH_4_Cl for 2 h and then incubated with or without f-Aβ_1-42_ and eHsp90α as in F. Anti-MAPI-LC3B antibodies were used to detect MAPI-LC3B expression in the immunoblots. Samples containing equal amounts of protein were loaded and equal loading was confirmed by the anti-GAPDH antibody immunoblot. Experiments were repeated at least three times, reproducibly.

We also observed increased MAP1-LC3B-II protein expression in BV-2 cells incubated with f-Aβ_1-42_ and eHsp90α, potentially suggesting increased autophagy under these conditions (Fig. 6F). Intracellular levels of MAP1-LC3B-II are generally correlated with the numbers of autophagosomal puncta [27]. When pretreated with lysosomal inhibitor ammonium chloride for 2 h, levels of MAPI-LC3B-II, (the autophagosome-associated, lipidated form of MAP1-LC3 that accumulates due to inhibition of lysosomal degradation), were increased in cells treated with both f-Aβ_1-42_ and eHsp90α (Fig. 6G).

### eHsp90α induces increased levels of autophagy proteins p62 /SQSTM1) and MAPI-LC3B in microglia

Among candidate intermediates in the autophagy pathway, MAPI-LC3B and p62 (encoded by *Sqstm1*) play particularly significant roles in the processing of ubiquitinylated proteins targeted for degradation [28]. We therefore next examined the expression of MAPI-LC3B and p62 after eHsp90α treatment in BV-2 cells (Fig. 7 A, C). As NRF2 has recently been linked to autophagy, we hypothesized that this factor might function as a regulator of p62 and LC3 expression to mediate the observed increases in indicators of autophagy and increased clearance of fAβ_1-42._ To test this possibility, *Nfe2l2* transcript levels were reduced by RNA interference (Fig. 7 A, C). Relative levels of NRF2 proteins in BV-2 cells were increased by eHsp90α treatment but not by exposure to f-Aβ_1-42_ (Fig. 7A). LC3II levels in these samples were increased by the eHsp90α treatments but not by f-Aβ_1-42_, and were reduced in samples transfected with *Nfe2l2* mRNA-targeting siRNA (Fig. 7A). f-Aβ_1-42_ treatment increased LC3I protein levels but not LC3II levels (Fig. 7A).

**Figure 7.**
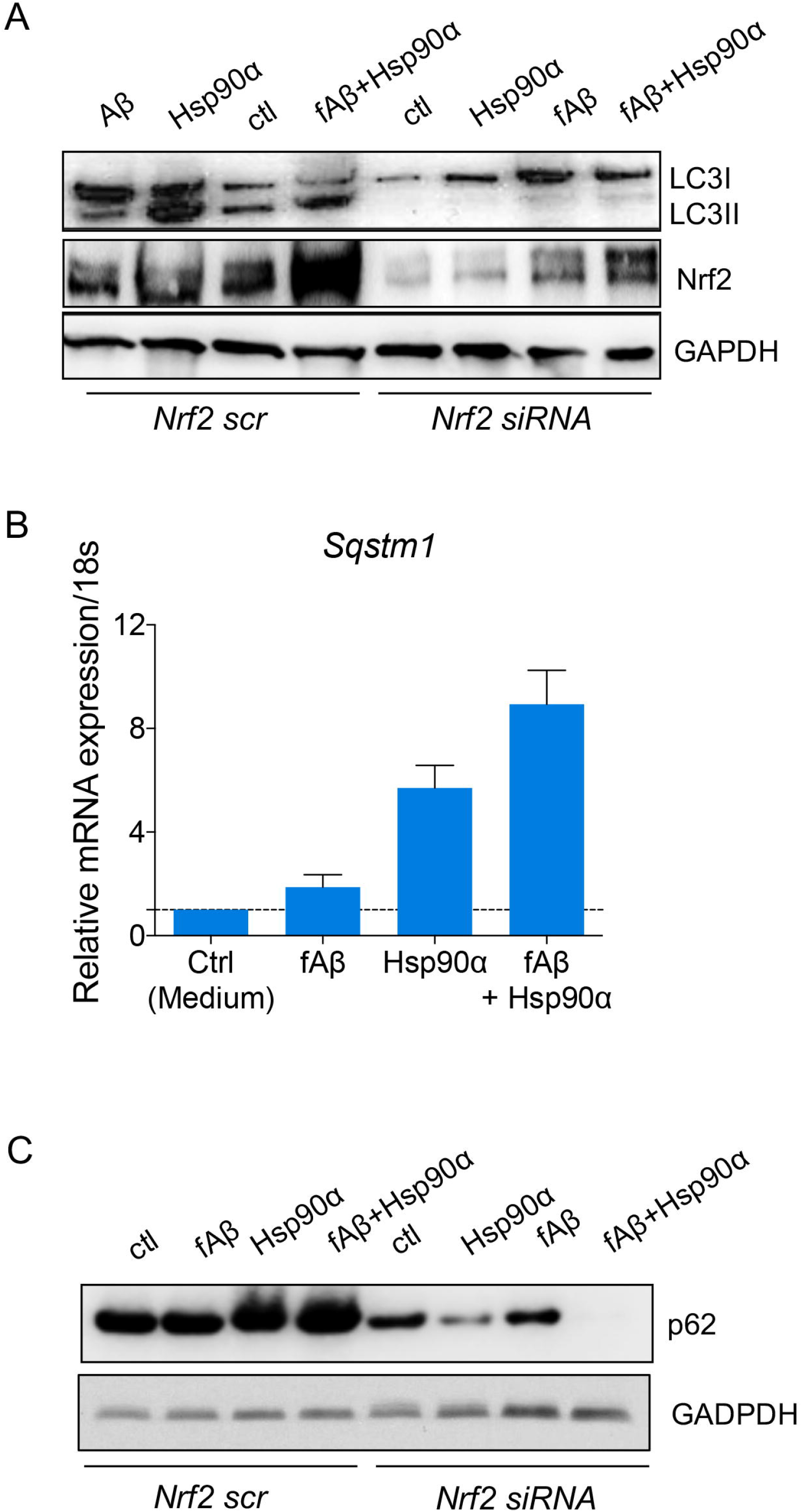
eHsp90α induced autophagy involves activation of NRF2 and subsequent expression of NRF2-regulated gene products. (A) BV-2 cells were transfected with either *scr* or *Nfe2l2* siRNA for 72 h. Cells were then incubated with f-Aβ_1-42_, eHsp90α and f-Aβ_1-42_ + eHsp90α for 4 h. Total cell lysates were collected and samples with equal amount of protein were separated using SDS-PAGE before transfer to PVDF membrane. Relative levels of the indicated proteins were determined by immunoblot using anti-MAPI-LC3B, and membranes then stripped and probed sequentially for NRF2 and GAPDH. (B) BV-2 cells were incubated with either vehicle control, f-Aβ_1-42_, eHsp90α ± f-Aβ_1-42_ or eHsp90α alone for 4 h. Total RNA was then extracted, and real-time qPCR was performed using primers to detect the *Sqstm1* transcript. *Sqstm1* mRNA levels were measured and represented relative to the amount of *18s* RNA. (C) BV-2 cells were transfected with either *Scr* or *Nfe2l2* siRNA for 72 h. Cells were then incubated with the indicated ligands for 4 h. Cell lysates were then collected and proteins separated by SDS-PAGE. p62 and GAPDH protein levels were detected sequentially by immunoblot using anti-p62 and anti-GAPDH antibodies. Experiments were performed three times with reproducibility.

At the mRNA level, eHsp90α treatment increased the levels of the p62 encoding mRNA of *Sqstm1* in BV-2 cells, with or without f-Aβ_1-42_ while exposure to the microfibrils alone had minimal effects (Fig. 7B). It might be significant that f-Aβ_1-42_ alone produced small increases in NRF2 and p62 mRNA, perhaps suggesting that the microglia were attempting to mount a homeostatic response to the toxic aggregates and that such a response could be amplified by eHsp90α treatment (Fig. 3A, 7B). To control for potential effects of NRF2 knockdown on f-Aβ_1-42_ uptake, BV-2 cells were transfected with *Nfe2l2*-targeting siRNA, treated with FITC-f-Aβ_1-42_ and analyzed by fluorescence microscopy after 72 h. We did not detect significant changes in uptake in cells depleted of NRF2 (Suppl. Fig. 4). These data therefore suggested that eHsp90α-mediated changes in f-Aβ_1-42_ uptake and entry into the autophagy pathways were independently mediated events. eHsp90α treatment also led to increased levels of p62 proteins with or without f-Aβ_1-42_ exposure, while f-Aβ_1-42_ alone produced minimal effects (Fig. 7C). eHsp90α-induced p62 was abrogated in cells transfected with *Nfe2l2*-targeting siRNA (Fig. 7C), again suggesting a primary role for this transcription factor in expression of autophagy intermediates.

## Discussion

The pathology of AD involves both intra- and extracellular Aβ deposition to neurons [29]. Intracellular Aβ plaques could be reduced by overexpression of HSPs such as Hsp40 and Hsp70, both of which bind to oligomers in the intra-neuronal space [30]. It has, however been a challenge to derive approaches to remove extracellular Aβ plaques from the brain, which are initially formed outside the nucleus and become neurotoxic [31]. Considering the chaperoning effects of Hsp90α and its role in internalizing antigens in immune cells, we hypothesized that eHsp90α might play a role in removing extracellular Aβ plaques in treatment of AD [7, 20].

Recent studies suggested a dominant role for microglia, in the progression of AD [32-34]. Microglia form a lineage originated from the yolk sac, which is distinct from other mononuclear phagocytes, and can play homeostatic roles in AD, by surrounding and removing Aβ plaques, apoptotic cells and debris [33, 34]. However, prolonged microglial activation may enhance the neurodegenerative phenotype of these cells in the pathology of AD, mediating neurotoxicity and loss of synapses and neurons [14]. Here we have demonstrated that the inflammatory neurotoxicity of f-Aβ_1-42_ exerted by microglia could be reduced in the presence of eHsp90α (Fig. 1). Our results suggested that eHsp90α may play a vital role both in engulfment and processing of extracellular f-Aβ_1-42_ aggregates (Fig.4). eHsp90α is internalized in a receptor-dependent pathway by phagocytic cells such as dendritic cells and macrophages [35, 36]. eHsp90α can bind to multiple receptors expressed in both non-immune cells and immune cells [37], including, but not limited to, LRP-1, SREC-I, LOX-1, and Feel-1/Stabilin-1 [6, 7, 36, 38, 39]. Other investigators have shown that non-fibrillar Aβ can be taken up by macrophages and microglia by receptor mediated phagocytosis [40]. Here we observed that eHsp90α causes enhanced uptake of f-Aβ_1-42_ in microglia (Fig. 5).

In addition to a method to recycle their own components during nutrient deprivation, cells utilize autophagy to process extracellular components such as dead cells and invading pathogens after phagocytosis [41, 42]. In this study, we demonstrated that the processing of extracellular f-Aβ_1-42_ by microglia is switched towards the autophagy pathway when Hsp90α is added to the extracellular media (Fig.6). It had been shown that the autophagy pathway is beneficial for AD, and that f-Aβ_1-42_ undergoes autophagy mediated processing in microglia [43]. However, microglial autophagy appears to be defective in some AD patients, a loss which may contribute to activation of microglia via the inflammasome [26]. We also observed that Hsp90α could increase MAPI-LC3B expression in microglia, and was found in MAPI-LC3B containing autophagosomes (Fig. 6). We observed a significant increase in MAPI-LC3B expression in cells with Hsp90α only and f-Aβ_1-42_ + Hsp90α, evidence that autophagy plays a role in Hsp90α mediated processing of internalized f-Aβ_1-42_ in microglia (Fig. 6). In autophagy, cytosolic MAPI-LC3B becomes conjugated to phosphatidylethanolamine, forming LC3BII on the autophagosomal membrane until this molecule fuses with lysosomes [44]. The other adaptor protein involved in the autophagy pathway, p62 / Sqstm1 is known to interact with LC3BII and facilitate ubiquitinylated protein targeting to autophagosomes and degradation [45]. We observed increases in both *LC3B* and *p62 /Sqstm1* mRNA and protein expression in the presence of eHsp90α suggesting the role of increases in these proteins in initiating this process (Fig. 7).

A central role in the response of microglia to Hsp90α appeared to be played by NRF2, which is induced and activated by the chaperone in microglia (Fig 3). Although NRF2 is classically activated by oxidative stress, such a mechanism seems unlikely here from our current understanding of the effects of extracellular Hsp90α. The mechanisms by which Hsp90α activate NRF2 are therefore unclear, but may involve downstream signaling cascades emanating from liganded HSP receptors [35]. Decreases in NO levels under conditions of activated NRF2 might be related to the known antagonism between this factor and NFkB, the key activator of inflammatory transcription [46]. Activated NRF2 can inhibit NFkB-mediated expression of *iNOS* as well as down-regulating inflammatory genes such as *COX-2, TNF*α and *IL1*β [46]. Recently it was shown that inflammatory responses induced by f-Aβ_1-42_ fibrils in microglia are decreased in these cells by induction of autophagy [47].

Our experiments therefore demonstrate that eHsp90α can modulate the processing of f-Aβ_1-42_ in microglia while mitigating the resulting oxidative burst. The mechanisms involved included induction of NRF2 and the oxidative stress response and recruitment of the autophagy pathway. These data may suggest potentially novel uses for HSPs in future treatment of AD.

## Acknowledgements

This research was supported by NIH research grants RO-1CA047407, RO-1CA119045 and RO-1CA094397 (for SKC) and RF1-AG054199 and R01-AG066429 (TI). A.M. is a recipient of a Harvard JCRT award and TJB was a recipient of a CAPES-PNPD fellowship. The authors alone are responsible for the content and writing of the paper. We thank the Department of Radiation Oncology, Beth Israel Deaconess Medical Center for interest and support.

^1^To whom correspondence should be addressed: Stuart K. Calderwood, Beth Israel Deaconess Medical Center, Harvard Medical School, 330 Brookline Ave, East Campus, DA-717A, Boston, MA 02215, USA.

## Data Availability

All data relevant to the manuscript are contained in the text figures and Supplementary data and any additional data will be made available.

## Notes

### Competing Interest Statement

The authors have declared no competing interest.

