## Supplemental Figures for "Extracellular Hsp90α Detoxifies β-Amyloid Fibrils Through an NRF2 and Autophagy Dependent Pathway"

### Slide 1
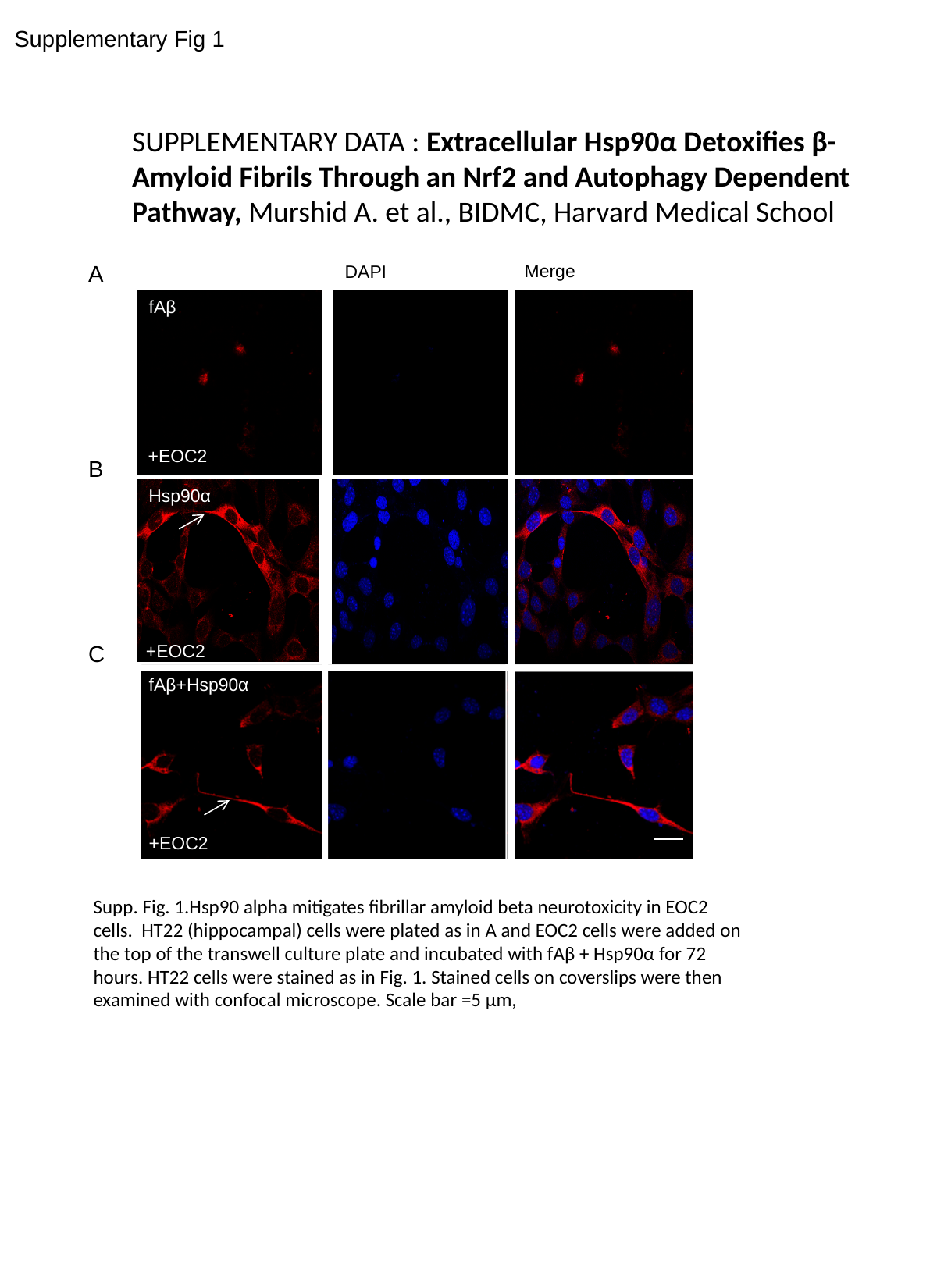

Supplementary Fig 1
SUPPLEMENTARY DATA : Extracellular Hsp90α Detoxifies β-
Amyloid Fibrils Through an Nrf2 and Autophagy Dependent
Pathway, Murshid A. et al., BIDMC, Harvard Medical School
A
Merge
DAPI
fAβ
+EOC2
B
Hsp90α
C
+EOC2
fAβ+Hsp90α
+EOC2
Supp. Fig. 1.Hsp90 alpha mitigates fibrillar amyloid beta neurotoxicity in EOC2 cells. HT22 (hippocampal) cells were plated as in A and EOC2 cells were added on the top of the transwell culture plate and incubated with fAβ + Hsp90α for 72 hours. HT22 cells were stained as in Fig. 1. Stained cells on coverslips were then examined with confocal microscope. Scale bar =5 µm,

### Slide 2
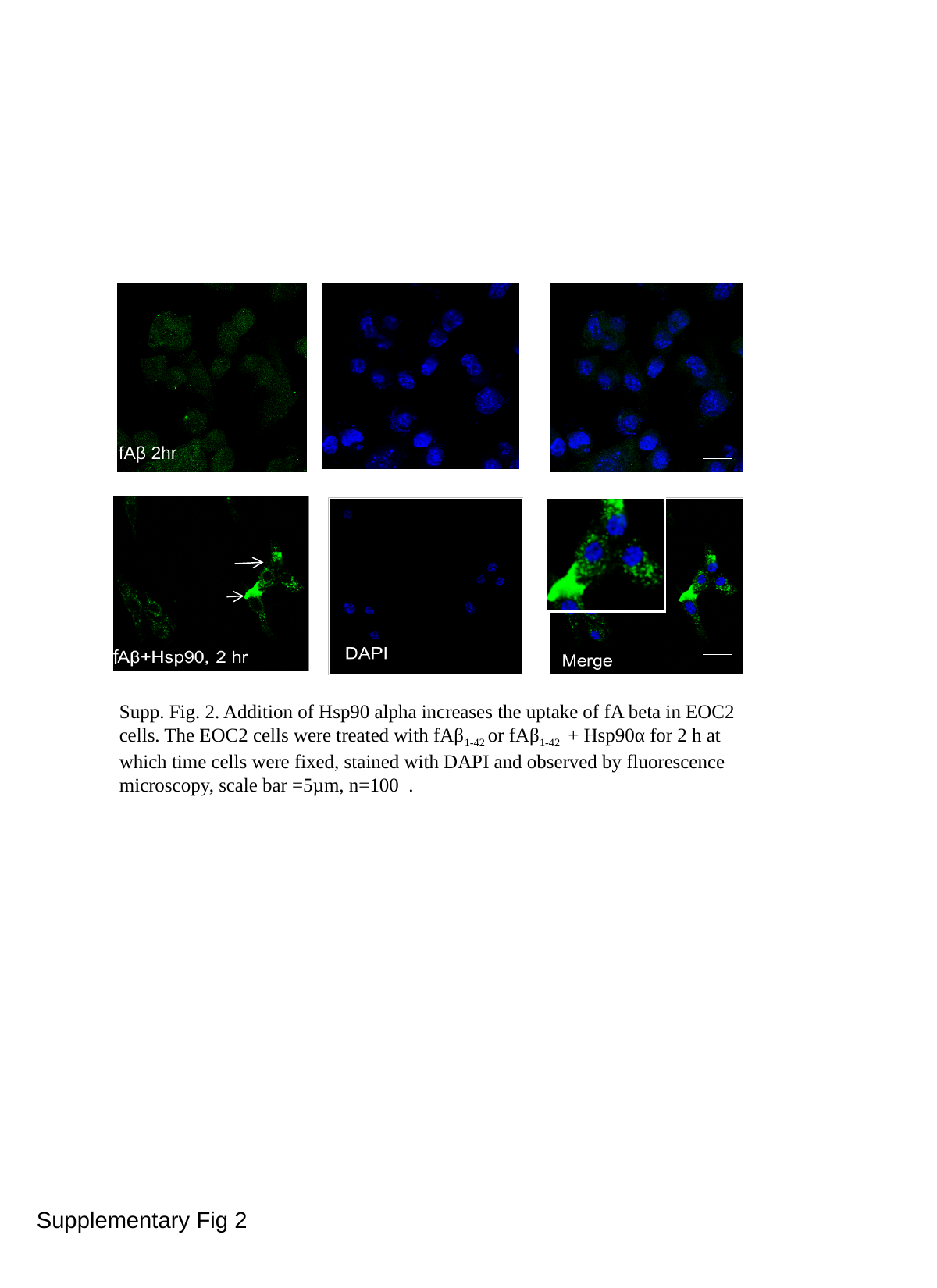

fAβ 2hr
f
Supp. Fig. 2. Addition of Hsp90 alpha increases the uptake of fA beta in EOC2 cells. The EOC2 cells were treated with fAβ_1-42_ or fAβ_1-42_ + Hsp90α for 2 h at which time cells were fixed, stained with DAPI and observed by fluorescence microscopy, scale bar =5µm, n=100 .
Supplementary Fig 2

### Slide 3
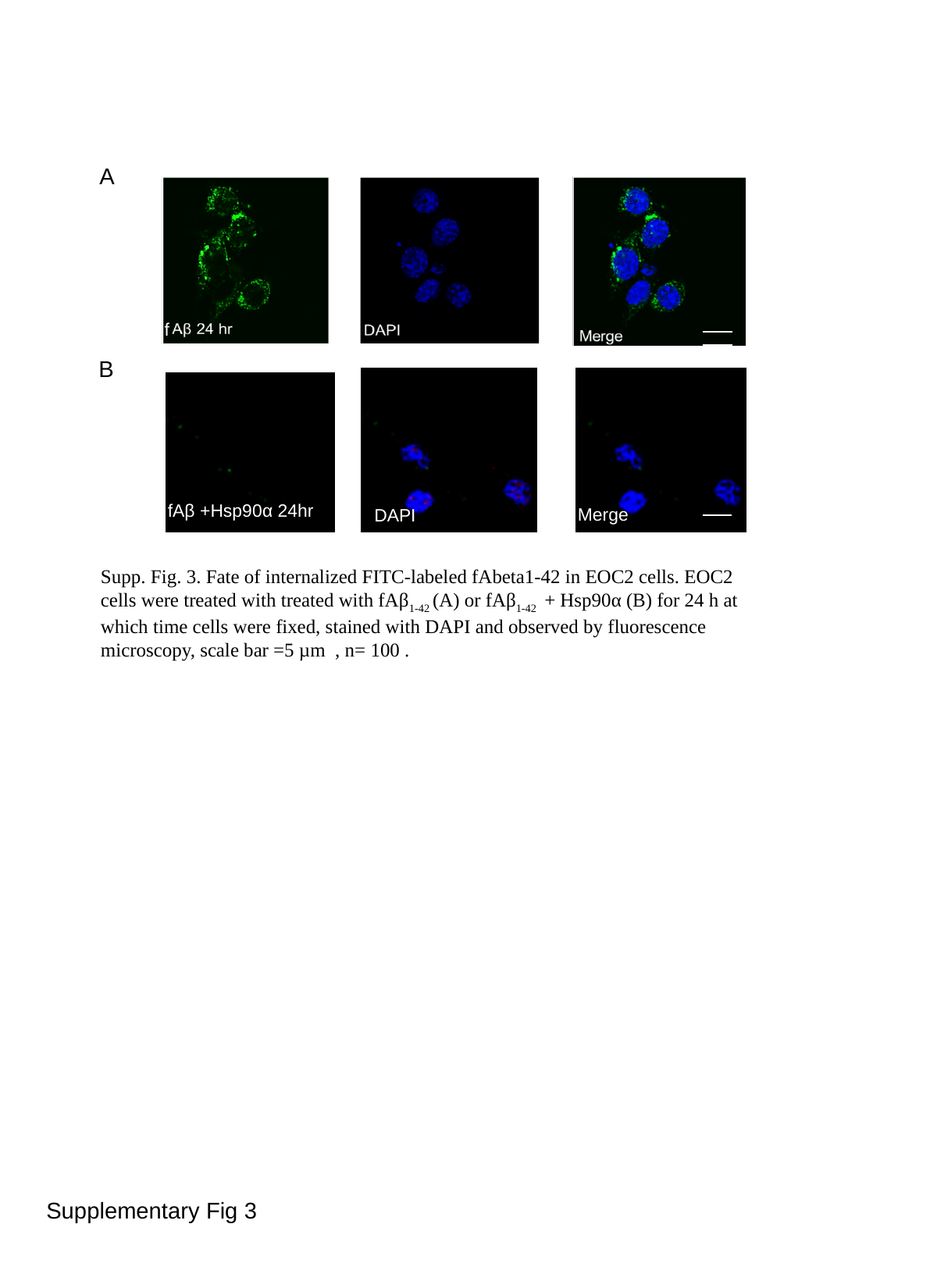

A
f
B
fAβ +Hsp90α 24hr
Merge
DAPI
Supp. Fig. 3. Fate of internalized FITC-labeled fAbeta1-42 in EOC2 cells. EOC2 cells were treated with treated with fAβ_1-42_ (A) or fAβ_1-42_ + Hsp90α (B) for 24 h at which time cells were fixed, stained with DAPI and observed by fluorescence microscopy, scale bar =5 µm , n= 100 .
f
Supplementary Fig 3

### Slide 4
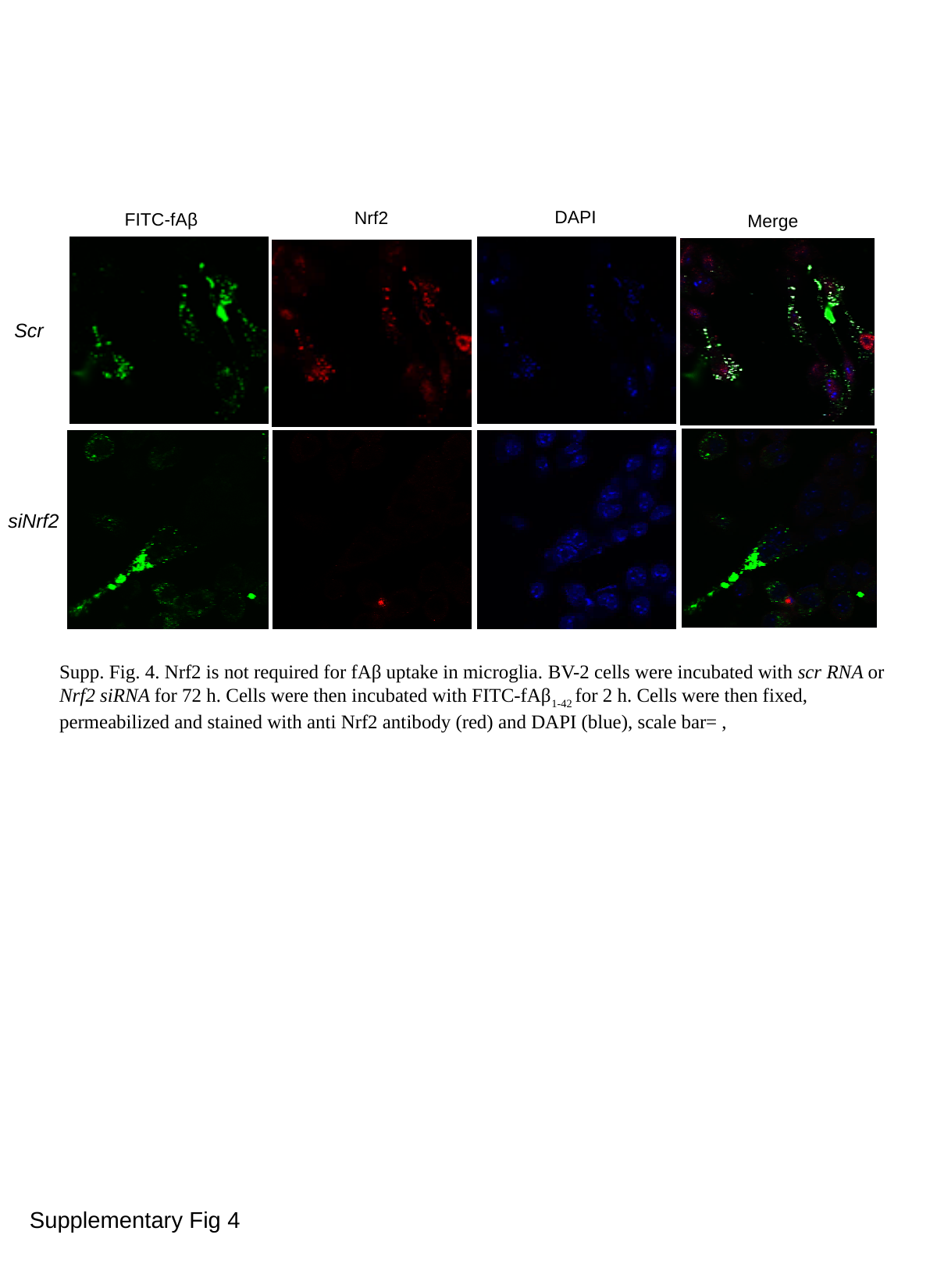

DAPI
Nrf2
FITC-fAβ
Merge
Scr
siNrf2
Supp. Fig. 4. Nrf2 is not required for fAβ uptake in microglia. BV-2 cells were incubated with scr RNA or Nrf2 siRNA for 72 h. Cells were then incubated with FITC-fAβ_1-42_ for 2 h. Cells were then fixed, permeabilized and stained with anti Nrf2 antibody (red) and DAPI (blue), scale bar= ,
Supplementary Fig 4
